# A novel tool for standardizing clinical data in a realism-based common data model

**DOI:** 10.1101/2020.05.12.091223

**Authors:** Hayden G. Freedman, Heather Williams, Mark A. Miller, David Birtwell, Danielle L. Mowery, Christian J. Stoeckert

## Abstract

Standardizing clinical information in a common data model is important for promoting interoperability and facilitating high quality research. Semantic Web technologies such as Resource Description Framework can be utilized to their full potential when a clinical data model accurately reflects the reality of the clinical situation it describes. To this end, the Open Biomedical Ontologies Foundry provides a set of ontologies that conform to the principles of realism and can be used to create a realism-based clinical data model. However, the challenge of programmatically defining such a model and loading data from disparate sources into the model has not been addressed by pre-existing software solutions. The PennTURBO Semantic Engine is a tool developed at the University of Pennsylvania that works in conjunction with data aggregation software to transform source-specific RDF data into a source-independent, realism-based data model. This system sources classes from an application ontology and specifically defines how instances of those classes may relate to each other. Additionally, the system defines and executes RDF data transformations by launching dynamically generated SPARQL update statements. The Semantic Engine was designed as a generalizable RDF data standardization tool, and is able to work with various data models and incoming data sources. Its human-readable configuration files can easily be shared between institutions, providing the basis for collaboration on a standard realism-based clinical data model.

## 1. Introduction

In biomedicine, significant challenges for biomedical researchers working with clinical data include 1) non-centralized, disparate data sources and 2) varying standards of knowledge representation between these data sources. A large hospital system may have various systems for capturing data, perhaps established at different times or with varying intent, and which are often not well integrated.^1^ Additionally, many hospital systems use unique and proprietary standards for storing clinical data, making data sharing between institutions difficult.^2^

Enabling the interoperability and exchange of clinical data is crucial for medical progress in the digital age.^3^ Common data models (CDMs) standardizing the storage of clinical data are available from the Observational Medical Outcomes Partnership (OMOP) ^4^, the National Patient-Centered Clinical Research Network (PCORnet)^5^, the Informatics for Integrating Biology & the Bedside (i2b2)^6^, and the Health Level 7 Fast Healthcare Interoperability Resources (HL7 FHIR)^7^.

However, none of these solutions leverages realism-based biomedical ontologies. The Open Biomedical Ontologies (OBO) Foundry^8^ provides a set of ontologies constructed using realism-based principles. In the seminal paper on ontological realism, Smith and Ceusters assert this fundamental idea:

> The most effective way to ensure mutual consistency of ontologies over time and to ensure that ontologies are maintained in such a way as to keep pace with advances in empirical research is to view ontologies as representations of the reality that is described by science.^9^

In addition to ensuring consistency and accuracy of ontologies, the realism-based philosophy can be advantageous for understanding clinical data. These data are only meaningful because they are about real patients and their situations, and a realism-based data model makes this explicit. In a recent paper, Bona et. al argue that semantic Web technologies such as Resource Description Framework (RDF) and Ontology Web Language (OWL) cannot be used to their full potential if shallow, one-to-one mappings between data elements and ontology classes are implemented. A semantically rich model can draw on classes provided by OBO Foundry ontologies to assist in effectively representing, retrieving, and axiomatically inferring implicit information about data.^10^ Additionally, data elements within such a model can be seamlessly integrated with associated domain knowledge.^11^

The **PennTURBO (Transforming and Unifying Research with Biomedical Ontologies)** Group at the University of Pennsylvania^12,13^ developed a realism-based model for representing clinical data. This involved constructing a set of ontology classes that describe the domain of interest and defining how instances of those classes may relate to each other. Rather than only creating one-to-one mappings between data elements and ontology terms, we instantiate classes even if they are not directly mapped to data elements. We strive to provide a more complete representation of the meaning of the clinical situation. This approach allows flexibility in querying the database and promotes ease of comprehension towards the goal of achieving a widely accepted realism-based model^10^.

Despite the perceived benefits of adopting realism-based principles for the representation of clinical information, we found it difficult to document our model and apply it to instance level data. There are commonly used tools to build and maintain ontologies, such as Protégé^14^. However, our data model is not an ontology, but rather a specification for storing collected data as instances of classes derived from ontologies. The lack of an appropriate software tool for building and sharing realism-based models is one possible explanation for why they have not been widely adopted. This tool would need to facilitate collaboration between ontologists who understand the model and programmers who understand the data sources. Even if such a tool were to exist, there is also no reliable software available to apply a realism-based model to a clinical dataset.

We faced three distinct challenges: 1) define our realism-based model in a format that supports consistency, pattern re-use, self-documentation, and self-validation, 2) develop an approach for automatically implementing the model against heterogeneous data sources, and 3) facilitate ease of adoption at other institutions and agile expansion as needed.

Our solution to these challenges is a software application called the *PennTURBO Semantic Engine* (from now on referred to as the *Semantic Engine*). The remainder of this paper will be devoted to introducing and demonstrating the utility and performance of this tool. First, we present related work. Then we describe the Semantic Engine and novel semantically driven methods for interpreting various data sources. Finally, we discuss the performance of the Semantic Engine over large, synthetic clinical datasets.

## 2. Background and Related Works

Data represented with the Resource Description Framework (RDF) are composed of triples using a three-part structure involving a subject, a predicate, and an object. Such a triple asserts a relationship between the subject and the object. RDF triplestores^15^ are graph databases designed for the storage and retrieval of triples through semantic queries. The query language for RDF is SPARQL, which functions by matching a given triple pattern against a triplestore. At most healthcare institutions, the majority of clinical data is stored within relational database tables, not in an RDF format.

A 2015 paper describes the development of a software tool with similar goals to ours. Mate et. al describe the need for transforming data from heterogeneous clinical sources and enabling data interoperability between institutions.^16^ Although they are not concerned with ontological realism, they determined that RDF-based ontologies would be useful for abstracting their source and target schemas as well as the mappings between them. They recognize that flexibility in adapting a configuration is achieved through modularity, and thus emphasize the separation of the source ontology from the source data and the mappings from the source and target ontologies. A major difference between our efforts is that their final output is a SQL database, where ours is an RDF repository.

We have observed from existing literature that the transformation of relational data into RDF has been attempted using both *single-step mapping* and *two-step mapping* approaches. A *single-step mapping* approach means transforming data from a relational schema into an RDF schema that mimics the original relational schema. A *two-step mapping* means transforming data from a relational schema into an intermediary RDF representation of that schema, then harmonizing it with a target model that does not depend on any other schema. The solution we present in this paper is the second step of a *two-step mapping* solution in which the target model is not dependent on the schemas of any other databases.

### 2.1 Single-Step Mapping Software from Relational Schemas to RDF

*R2RML (RDB to RDF Mapping Language)*^17^ is a relational database to RDF mapping language which uses an RDF-based syntax. It is incorporated into a World Wide Web Consortium (W3C)^18^ standard. R2RML facilitates generating a dump of RDF data or querying a virtual SPARQL endpoint, a volatile, in-memory reshaping of what is durably stored in a relational format.

Several tools have been built on top of the R2RML framework, such as *Karma*^*19*^, a software tool that can be used to create single-step mappings between a relational data source and an RDF model.^20^ Developed by Knoblock et al. at USC’s Center on Knowledge Graphs, Karma provides a visual interface and allows the user to work directly with a subset of the incoming data, along with ontology terms or axioms. The interface allows a user to “drag and drop” to map ontology classes to relational table columns and to define relationships between columns.

Another tool that relies on an implementation of R2RML is *Ontop*^*21*^, a system that exposes the content of relational databases as RDF. While many tools in this realm provide a set of RDF triples based on the content of a relational database, Ontop differs in that it queries the original database directly by translating SPARQL queries into SQL queries. Ontop is available as a plugin for the ontology editor Protégé to edit and test mappings.

Bernabé-Díaz et al. created the *Semantic Web Integration Tool (SWIT)*^22^ at the TECNOMOD group of the Department of Informatics and Systems from the University of Murcia. SWIT is a tool that defines single-step mappings from a relational schema to an RDF model.^23^ This system uses XML configuration files and a web interface to create mappings.

### 2.2 Two-Step Mapping Software from Relational Schemas to RDF

Sun et al. use application ontologies to initially create RDF data from relational sources, and then use SPARQL templates and the EYE reasoner to complete the transformation to a source-independent RDF model based on a canonical ontology. Their primary goal is to enable the export of data from their canonical ontology model into formats accepted by various other systems^24^.

Berges et al. describe an ontology-based approach to normalizing and synchronizing clinical data that uses a two-step mapping. They apply a source-independent RDF model built from a canonical ontology with terms that correspond to codes in pre-existing medical terminologies. A main focus is on smoothing out abnormalities imposed by aggregating data from several heterogeneous relational systems. They map their application ontology to their canonical ontology by defining axioms and using a SWRL reasoner^25^.

Birtwell et al. developed *Carnival*^26^ at the University of Pennsylvania. Carnival is an open-source data integration and aggregation application for clinical data, which uses a set of data source adapters to pull data from various sources, including relational databases, flat files, and internal APIs, into a property graph. The Carnival property graph leverages a core schema inspired by the Ontology for Biobanking (OBIB)^27^ and the Ontology for Biomedical Investigations (OBI)^28^. Carnival writes concise RDF data from the property graph using a set of data writers. The codebase is modular and can be modified to integrate new data sources. We use Carnival as the first step of our two-step solution, though other tools can also be used.

Given these aforementioned software tools and approaches, we observed several opportunities for innovation. First, the single-step mapping solutions rely on approaches in which the target model is not defined separately from the mappings to a data source. This leads to difficulty in comprehending the target model, misses the opportunity for data validation prior to executing the transformation, and does not facilitate the standardization of data between heterogeneous sources. Second, although several of the two-step approaches might partially address the concerns of a single-step mapping solution, these solutions do not create a target model with the semantic richness to reflect the true clinical reality.

### 2.3 RDF Validation Languages

Our system uses a custom approach for the validation of input data that is described in **Section 3.7**. Shapes Constraint Language (SHACL)^29^ and Shape Expressions (ShEx)^30^ are two alternative languages which enable validation of RDF instance data in accordance with a provided specification. These two languages have similar goals and share some common features; however, they have continued to evolve separately. SHACL is concerned with defining constraints and implementing checks against RDF instance data. ShEx is used for RDF graph validation as well, but was originally intended to also act as a schema for RDF graphs.^31^ At the time of building our system, neither SHACL nor ShEx was supported by our triplestore.

### 2.4 A Novel Semantically Driven Transformation of Clinical Data

In this manuscript, we propose a novel solution, the **PennTURBO Semantic Engine**, an open-source, RDF data transformation tool developed with realism-based principles. The Semantic Engine uses one set of transformation instructions per data source to transform clinical data into a realism-based target model in a transparent and easily validated way. By keeping the specification of the realism-based model separate from the transformation instructions for a given data source, our system permits an ontologist to create a source-independent model, which can then be entirely or partially implemented for a given data source by a programmer.

The Semantic Engine uses an RDF-based domain-specific language (DSL) called the Semantic Engine Language, which rigorously defines the realism-based model and a set of transformation instructions for each data source. The Semantic Engine Language uses a relationship-oriented syntax that facilitates reliable and straightforward modifications or additions, using the SPARQL language as its underlying implementation. Rather than necessitating that a user develop proficiency with the SPARQL language, the Semantic Engine Language provides a direct interface for comprehending and manipulating the model.

Additionally, the Semantic Engine Language:

- ensures that transformation instruction modules build an interpretable subset of the realism-based model
- provides a single point-of-access for all relevant information for a given graph relationship, facilitating convenient re-usage
- ensures that similar data coming from different sources will be represented identically
- facilitates validation of incoming RDF triples against a given set of transformation instructions
- provides human-readable documentation of the realism-based model.

**Figure 1** shows a high-level overview of how ontologists and programmers can collaborate on the transformation of heterogeneous data into a realism-based model using components of the Semantic Engine. In the next section, we utilize a minimal example to describe the process of building a Semantic Engine configuration and executing it to transform data from a relational source.

**Figure 1:**
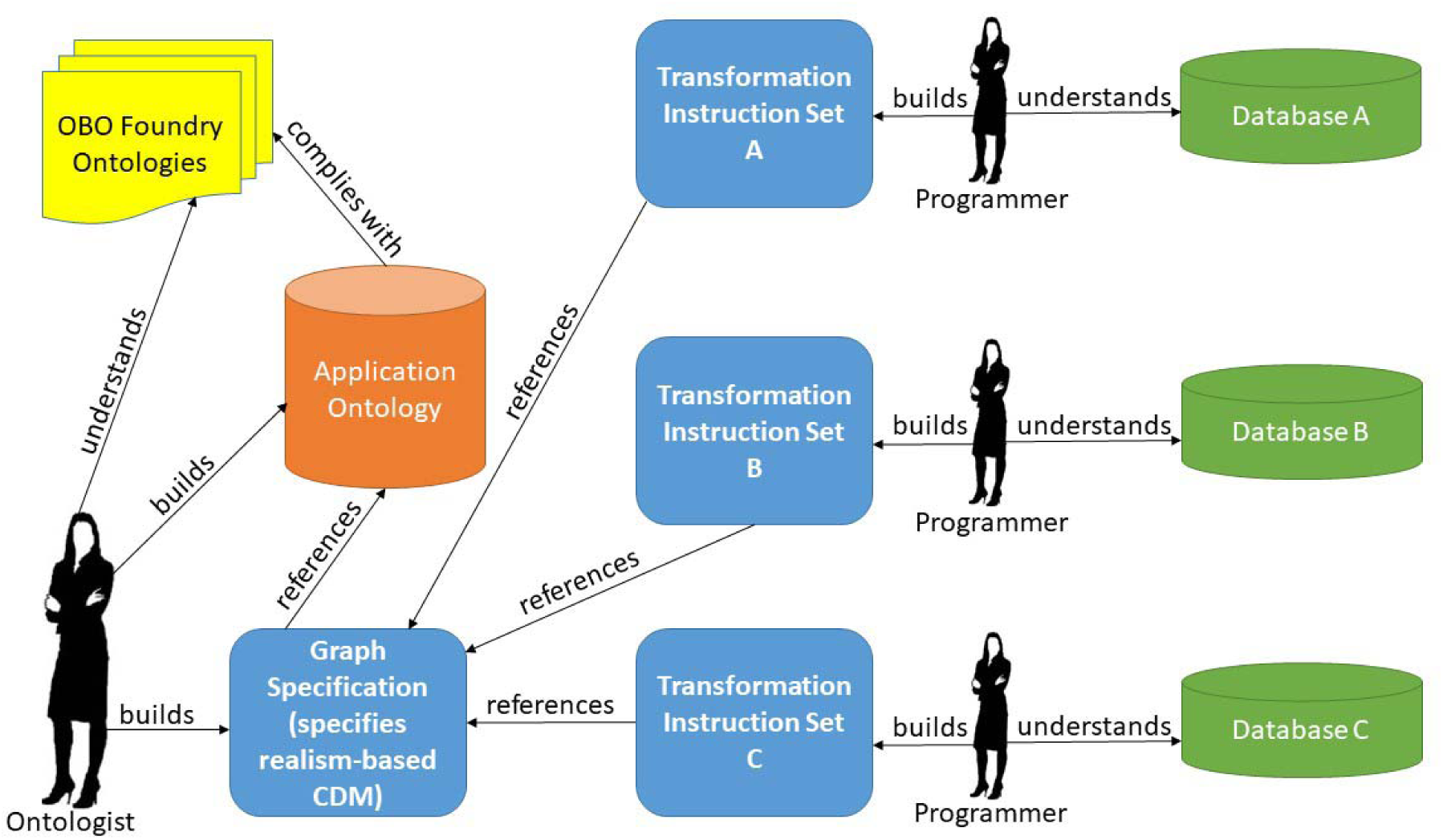
Workflow for building a Semantic Engine configuration. The objects colored in blue are coded using the Semantic Engine Language.

## 3. Semantic Engine Overview - Database Example

Let’s assume that we have a relational database, which we will call “A”, which stores only the most minimal of patient information: a single column of person identifiers. **Table 1** represents Database A.

**Table 1:**
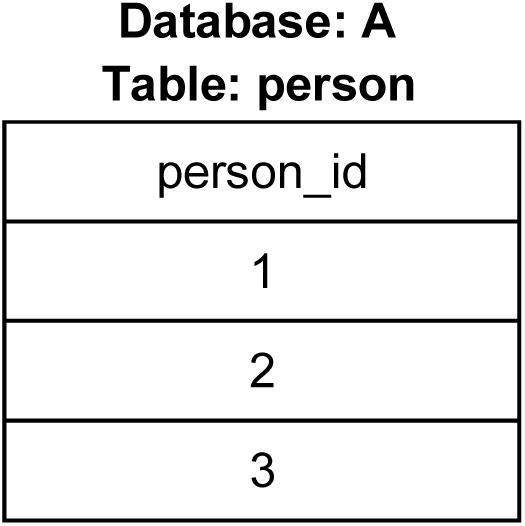
Sample Database A holding patient identifiers in column “person_id”

Our goal is to transform this relational data into a representation using our realism-based model. The Semantic Engine Language allows us to apply only the subset of our model which is relevant for a given data source. In this section, we will explore the architecture of the Semantic Engine and the syntax of the Semantic Engine Language by stepping through an example of how the Semantic Engine can be configured to accept data from Database A.

### Step 1: Convert the relational data in Database A into concise RDF triples

We are going to use a simple RDF schema and directly map^32^ the relational data to RDF. For each patient identifier present in Database A, instances of **inputSchema:homoSapiens** are created and linked to the literal identifier using the **inputSchema:identifier** predicate. The triples in **Figure 2** are then inserted into our RDF triplestore.

**Figure 2:**
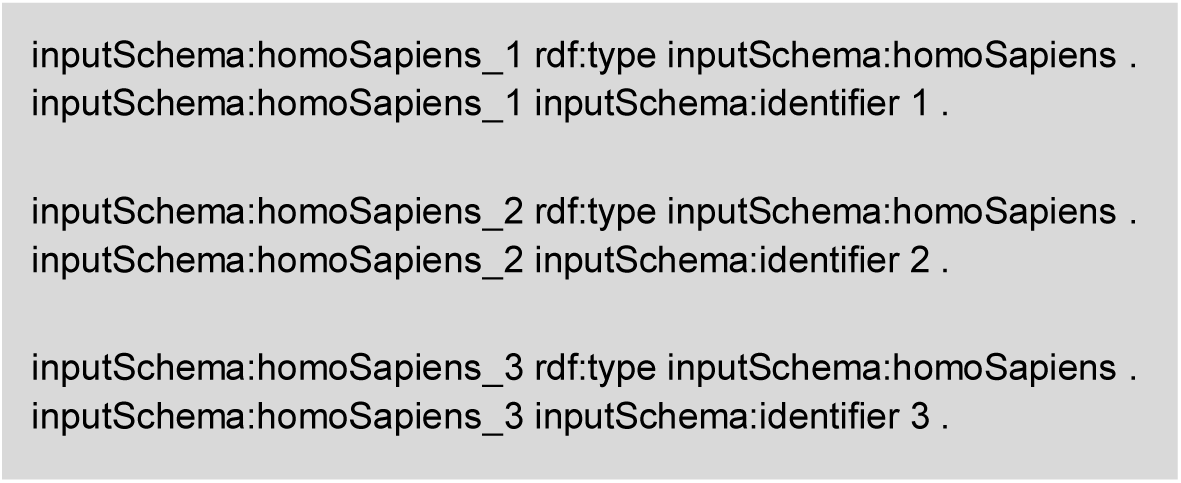
Source-specific triples mapped from Database A as input to Semantic Engine. “rdf:type” is the instantiation relation, which asserts that the subject (on the left) is an instance of the object (on the right).

At this point, the Semantic Engine has access to the data as concise triples. However, we will need to define mappings so that the software can understand data from this input source. That’s where the Semantic Engine Language comes into play.

### Step 2: Interpret concise RDF triples using Semantic Engine

The Semantic Engine Language configuration for a data source involves a collection of four files: an *application ontology*, a *Graph Specification*, a *Transformation Instruction Set*, and the *Semantic Engine Language ontology*. The application ontology, Graph Specification, and Semantic Engine Language ontology files are data source-agnostic, while the Transformation Instruction Set file is data source-specific. **Figure 3** shows a high-level overview of the components of the Semantic Engine and how they interact with various external components and data. In the following subsections, we will look at these components in detail in the context of defining the transformation for Database A.

**Figure 3:**
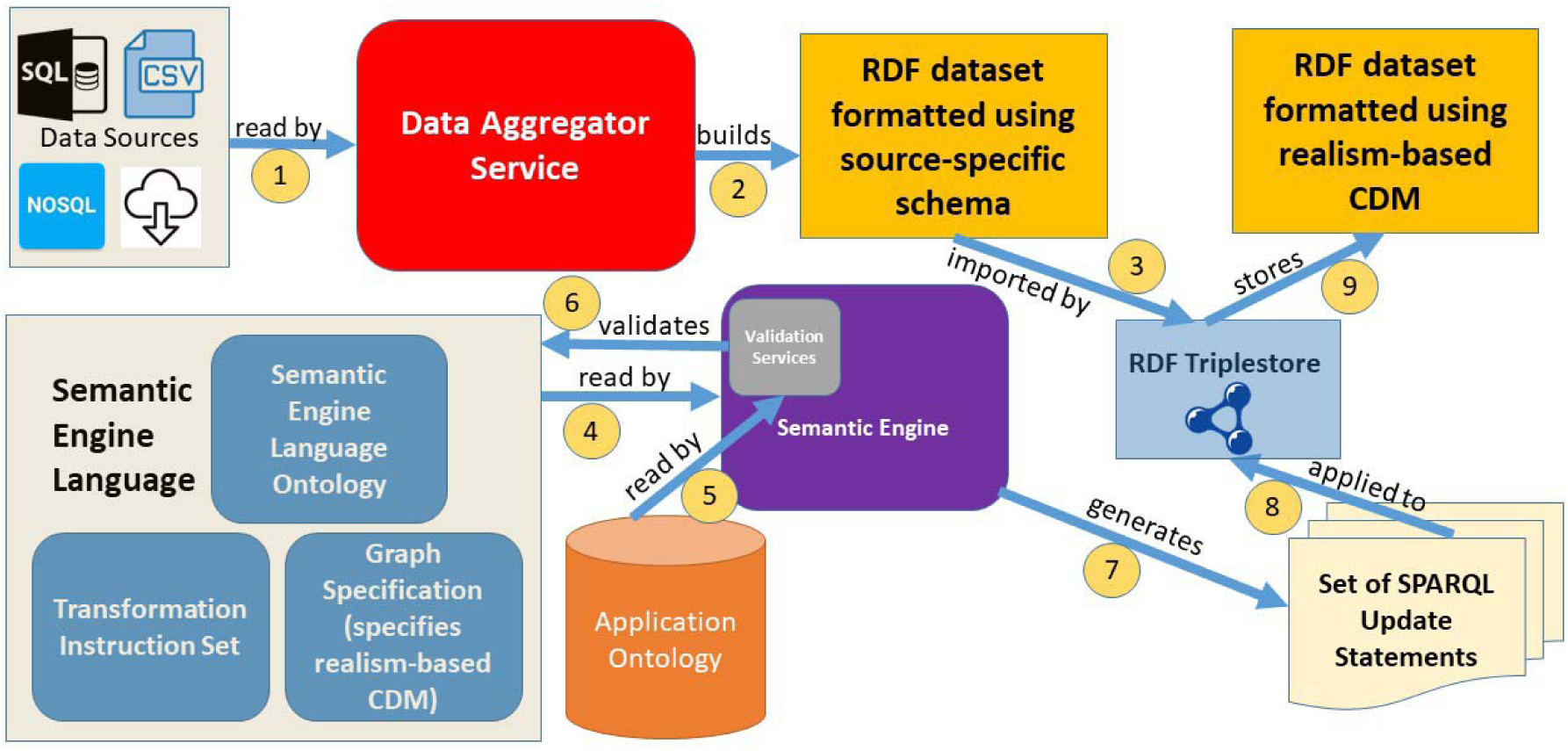
High-level overview of the components of the Semantic Engine and their interactions with various data sources to build a pipeline

#### 3.1 Application Ontology

##### Define all classes and predicates using realism principles

An application ontology is an ontology designed for a specific use, but which may import classes from more general ontologies. It may also provide additional native classes and predicates covering the cases in which no term exists in other ontologies to describe a relevant concept. The Semantic Engine application ontology file defines all of the classes and predicates which may be used in the realism-based model. The application ontology is the main point of reference for validating any implementation of the realism-based model. The first step in creating additions to the realism-based model would be deciding which terms are relevant to add to the application ontology.

#### 3.2 Semantic Engine Language - Graph Specification

##### Define relationships between instances in the realism-based model

The Graph Specification file is where the realism-based model is fully specified. It references classes and predicates from the application ontology. Unlike the application ontology, it defines all the ways that instances of particular classes may be connected with instances of other classes, classes themselves, or literal values. By making explicit the relationships by which two entities may be connected, it provides a representation of the realism-based model independent of any data source or set of transformation instructions. Another way to understand the role of this component is that it provides a set of constraints on the application ontology.

The Graph Specification is composed using the syntax of the Semantic Engine Language. The Semantic Engine Language represents relationships as instances of RDF objects called Connection Recipes, which will be discussed further in Section 3.6. Each Connection Recipe is specified with a Uniform Resource Identifier (URI). **Figure 4** is a subset of a possible Graph Specification file, showing only the Connection Recipes that relate to patient identifier data. The classes come from the PennTURBO Ontology^33^, the application ontology our group has developed. Classes in this figure are annotated with comments including the human-readable class label.

**Figure 4:**
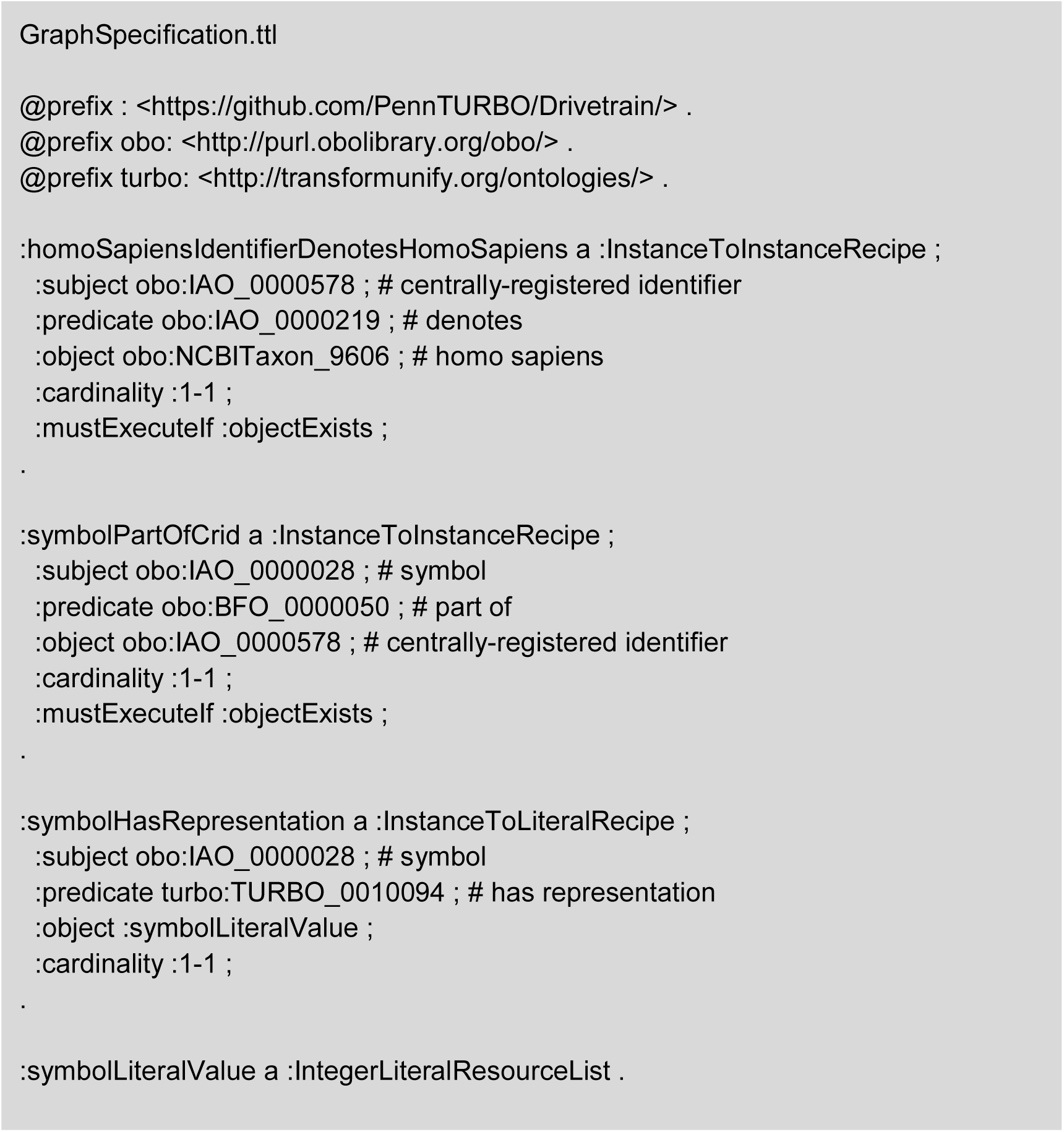
A subset of the Graph Specification showing only Connection Recipes specifically relevant to the data in Database A

In order to specify that the existence in reality of a person is denoted by an identifier in our database, we instantiate an instance of a homo sapiens only when there is an instance of a patient identifier. Therefore, it would not be appropriate to have an instance of Homo sapiens without an associated instance of a patient identifier. This concept is represented by the following triple in the Graph Specification:

:homoSapiensIdentifierDenotesHomoSapiens :mustExecuteIf :objectExists.

This statement references the Connection Recipe

**:homoSapiensIdentifierDenotesHomoSapiens**, and tells us that if the object of that recipe (an instance of class homo sapiens) exists, the subject of that recipe (an instance of class centrally-registered identifier, or CRID) must also exist, and be attached to the object by the predicate of that recipe, “denotes”. The ability to define rules for instantiating instances of classes based on the presence of specific incoming data fields was a key requirement for implementing the realism-based model. If at any point an instance of class Homo sapiens existed without an associated instance of a CRID, it would be reported by the validation service, which will be discussed further along in this section.

Connection Recipes can represent general patterns that can be re-used within the model. For example, recipes **:symbolPartOfCrid** and **:symbolHasRepresentation** are not specific to CRIDs or symbols associated with people and can be used to help instantiate any type of CRID. Since the same Recipe can be referenced multiple times, it would be trivial to implement, say, CRIDs and symbols denoting a prescription or a diagnosis by referring to the same **:symbolPartOfCrid** and **:symbolHasRepresentation** recipes. This re-usability encourages consistency in pattern development and helps reduce errors introduced by duplicating code.

#### 3.3 Semantic Engine Language - Transformation Instruction Sets

##### Define the transformation of source-specific triples to instantiate a partial realism-based model

While the Graph Specification provides a set of constraints on the application ontology but has no knowledge of incoming data sources, the Transformation Instruction Set maps concise source-specific triples (like in **Figure 2**) to a subset of the realism-based model as defined in the Graph Specification.

A new Transformation Instruction Set must be created for each new data source that is incorporated. Like the Graph Specification, Transformation Instruction Set files are composed with the syntax of the Semantic Engine Language.

The process of making a new Transformation Instruction Set based on source-specific RDF triples from a previously unincorporated data source first involves surveying the Graph Specification, in order to get a sense of how various relationships in the realism-based model fit together. Once a programmer has determined the subset of the Graph Specification that is relevant to their data source, they can begin writing new Connection Recipes in their Transformation Instruction Set that represent the source-specific RDF, and map them to pre-existing Recipes in the Graph Specification.

**Figure 5** shows what a Transformation Instruction Set for Database A could look like. It introduces a new type of RDF object, an Update Specification, which defines when and how a Connection Recipe will be executed.

**Figure 5:**
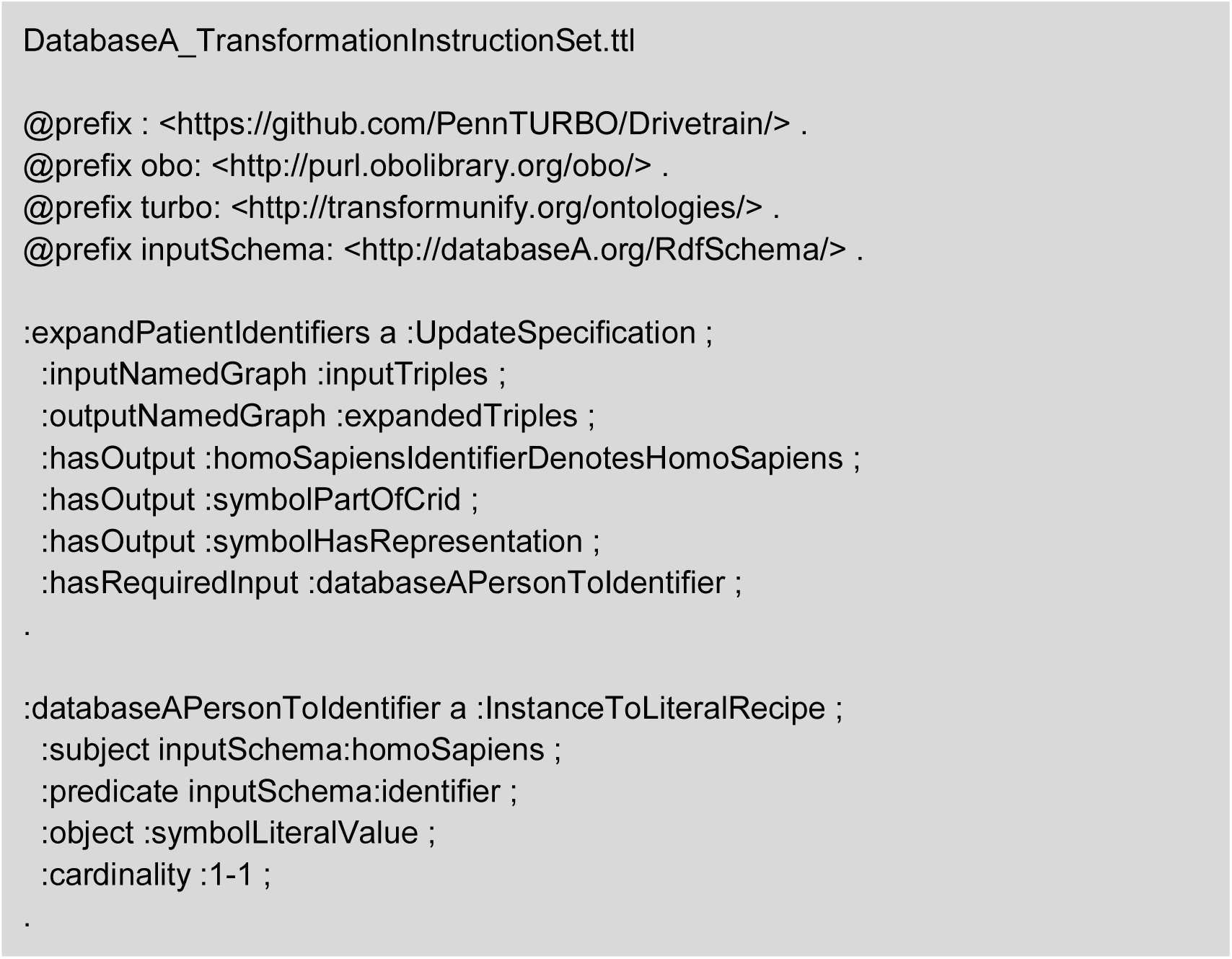
A Transformation Instruction Set for Database A

A valid Transformation Instruction Set must contain at least one instance of an Update Specification that defines input and output Connection Recipes. For incoming data sets that contain many fields, we build several Update Specifications, with each one transforming a specific subsection of the data. Since a single Connection Recipe could be the output of one Update Specification and the input to another, order matters. Therefore, it is possible to chain Update Specifications together within the Transformation Instruction Set.

#### 3.4 Semantic Engine Language Ontology

##### Define the classes, predicates, and associated terms for the application ontology

The Semantic Engine Language ontology file defines all of the classes and predicates available in the Semantic Engine Language. Following OBO Foundry guidelines, all classes defined in the TURBO application ontology are represented by a partially numeric URI. Our goal was to make using the Semantic Engine Language an intuitive experience for programmers and ontologists defining their own transformations, so the URIs of its classes and predicates are text-based and suggest the term’s function.

#### 3.5 Running the Semantic Engine

##### Step 3: Run the Semantic Engine using aforementioned configuration files

The Semantic Engine is written in Scala^34^ and can be run either as a precompiled .jar file or from the console of SBT, the build tool for Scala. Upon running the application, the Semantic Engine Language configuration files will be loaded into an RDF repository. Validation checks specified in the application code will ensure that the input data and Semantic Engine Language components are valid and consistent with each other. If these checks pass, the generated SPARQL updates will be executed in sequence against the input data using the RDF4J library^35^. The application will generate SPARQL update queries based on the contents of the Transformation Instruction Set.

**Figure 6** shows the SPARQL update that will be generated by running the Semantic Engine using the Graph Specification defined in **Figure 4** and the Transformation Instruction Set defined in **Figure 5**. SPARQL updates generated by the Semantic Engine have 3 distinct sections: INSERT, WHERE, and BIND.

**Figure 6:**
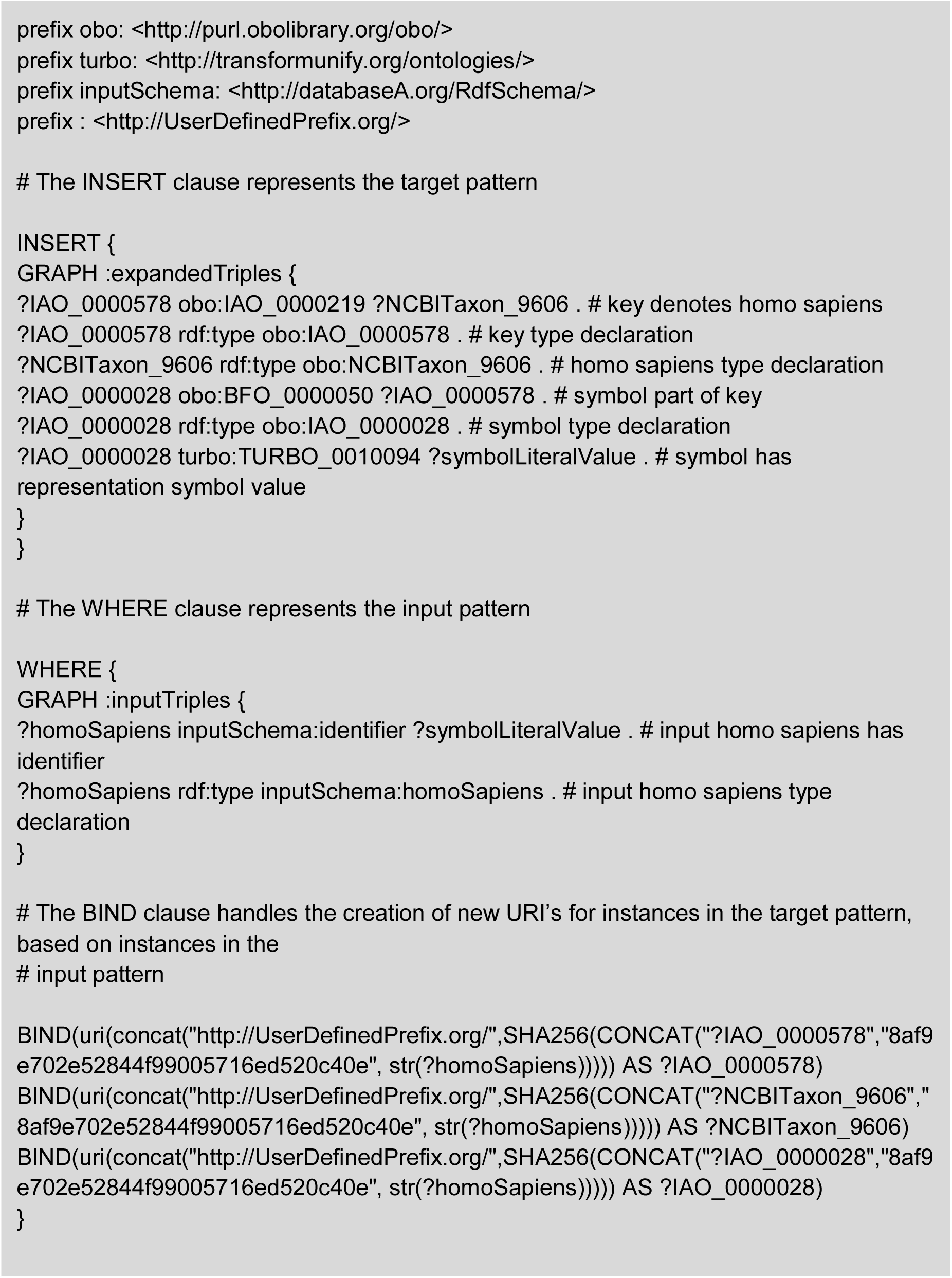
The SPARQL update generated by the Semantic Engine’s processing of Update Specification :expandPatientIdentifiers from Figure 5

1. The WHERE clause matches and binds variables against the source-specific RDF triples seen in **Figure 2**, using the pattern defined in **Figure 5**.
2. The BIND clause handles the creation of URIs for newly created instances. We create these URIs with the output of an SHA256 function which takes three parameters:

- a string representation of the type of the instance to be created, to ensure that new elements of different types will not have the same URI
- a string key specified by the user which acts as a “salt”, ensuring that the created URIs will be unique to this instantiation, unless the same key is used again
- a string representation of the URI identifying the instance with which the new instance should share a cardinality relationship (i.e. for each input of **inputSchema:homoSapiens**, we should create one new URI identifying an instance of type **obo:NCBITaxon_9606**, the OBO Foundry term for homo sapiens)
3. The INSERT clause assigns the bound variables to the pattern derived from the relevant subset of the Graph Specification in **Figure 4**.

#### 3.6 Diving Further into Connection Recipes

##### 3.6.1 Connection Recipe Overview

A Connection Recipe must include a subject, predicate, object, and cardinality declaration, and optionally specifies the conditions under which it must be executed. The predicate referenced by a Connection Recipe must be present in the application ontology. The subject may be present in the application ontology or it may reference a URI from an external vocabulary. The object can be present in the application ontology, reference an external URI, or represent a literal value.

Below are the six types of Connection Recipes, two of which are highlighted for their use in our minimal example.

1. *Instance-to-instance recipes* define a triple where the subject is an instance of a class and the object is also an instance of a class (see **:homoSapiensIdentifierDenotesHomoSapiens** in **Figure 4**)
2. *Instance-to-term recipes* define a triple where the subject is an instance of a class and the object is a class itself. This is used in the case that an instance has some direct relationship with a term in the associated application ontology, or to a term from an outside terminology.
3. *Term-to-instance recipes* are similar to Instance-to-term recipes, but the subject is the class and the object is the instance.
4. *Instance-to-literal recipes* define a triple where the subject is an instance of a class and the object is a literal value. The type of literal value may be further specified as instances of one of five literal types: string, Boolean, integer, double, or date. (see **:symbolHasRepresentation** in **Figure 4**)
5. *Term-to-literal recipes* define a triple where the subject is a class and the object is a literal value. We have used this type of recipe thus far mainly for retrieving classes from our application ontology that are linked to OMOP concept codes that are present in our input dataset.
6. *Term-to-term recipes* define a triple where the subject and the object are both classes. We have used this type of recipe thus far mainly for specifying subclass relationships.

The cardinality field of a Connection Recipe, specified by predicate **:cardinality**, defines how many instances of the subject may connect to an instance of the object via the stated predicate, and vice versa. The conditions that can be specified are **:1-1, :many-1**, and **1-many**.

The implementation requirement field, specified by predicate **:mustExecutetIf**, defines the conditions under which a given Connection Recipe must be implemented by a Transformation Instruction Set. There are several conditions that can be specified, including

**:objectExists** which we saw in **Figure 4**, as well as **:subjectExists** and

**:subjectOrObjectExists**.

##### 3.6.2 Representation in SPARQL

The SPARQL representation of a Connection Recipe depends on the type of entities that the recipe represents. Subjects or objects of recipes representing instances of classes will be automatically converted into a unique SPARQL variable based on the URI of the class. Types of instances are automatically assigned. **Table 2** demonstrates how the predicate used to link an Update Specification to a Connection Recipe affects where that Recipe will be placed in the generated SPARQL clause. **Table 3** contains examples of each of the six types of Connection Recipes as well as their subsequent compilation into SPARQL string snippets by the Semantic Engine.

**Table 2:** The predicate that links Connection Recipes to an Update Specification determines where the Connection Recipe will be placed in the generated SPARQL update statement

| Relationship between Update Specification and Connection Recipe | SPARQL Section |
| --- | --- |
| :hasRequiredInput | WHERE |
| :hasOptionalInput | WHERE (IN OPTIONAL BLOCK) |
| :hasOutput | INSERT |
| :removedBy | DELETE |

**Table 3:** Connection Recipes and their translation to SPARQL

| Connection Recipe Type | Example Representation in Semantic Engine Language Configuration | Example Representation in SPARQL |
| --- | --- | --- |
| *Instance-to-Instance* | :homoSapiensHasQualityBioSex a :InstanceToInstanceConnectionRecipe ;<br>:subject :homoSapiens ;<br>:predicate :hasQuality ;<br>:object :biologicalSex ;<br>:cardinality :1-1 ;<br>. | ?homoSapiens :hasQuality ?biologicalSex .<br>?homoSapiens rdf:type :homoSapiens .<br>?biologicalSex rdf:type :biologicalSex . |
| *Instance-to-Term* | :keyHasPartRegistryDenoter<br>a :InstanceToTermRecipe ;<br>:subject :key ;<br>:predicate :hasPart ;<br>:object :someRegistry ;<br>:cardinality :1-1 ;<br>. | ?key :hasPart<br>:someRegistry .<br>?key rdf:type :key . |
| *Term-to-Instance* | :registryDenoterPartOfKey<br>a :TermToInstanceRecipe ;<br>:subject :someRegistry ;<br>:predicate :partOf ;<br>:object :key ;<br>:cardinality :1-1 ;<br>. | :someRegistry :partOf<br>?key .<br>?key rdf:type :key . |
| *Instance-to-Literal* | :cridSymbolHasRepresentation<br>a :InstanceToLiteralRecipe ;<br>:subject :cridSymbol ;<br>:predicate :hasRepresentation ;<br>:object :tumor_LiteralValue ;<br>:cardinality :1-1 ;<br>. | ?cridSymbol<br>:hasRepresentation<br>?tumor_LiteralValue .<br>?cridSymbol rdf:type<br>:cridSymbol . |
| *Term-to-Literal* | :datumHasOmopConceptId<br>a :TermToLiteralRecipe ;<br>:subject :datum ;<br>:predicate :hasOmopConceptId ;<br>:object :omop_concept_id ;<br>:cardinality :1-1 ;<br>. | :datum<br>:hasOmopConceptId<br>?omop_concept_id . |
| *Term-to-Term* | :genderIdentityDatumSubclassOfDatum<br>a :TermToTermRecipe ;<br>:subject :genderIdentityDatum ;<br>:predicate rdfs:subClassOf ;<br>:object :datum ;<br>:cardinality :1-1 ;<br>. | :genderIdentityDatum<br>rdfs:subClassOf :datum . |

#### 3.7 Validation Services

The Semantic Engine software includes several layers of validation checks, broken up into modules called Validation Services. The overall objectives of the Validation Services are to ensure that all terms in the Semantic Engine Language are used as expected, ensure that created graph patterns are consistent both in the formal logical sense and with regards to the definition of the realism-based model, and ensure that the incoming source-specific RDF data matches the patterns specified by the Transformation Instruction Set. We break down the existing services below, in the order that the Semantic Engine executes them.

##### 3.7.1 Graph Specification Validation Service

The application ontology file defines all of the classes and predicates which may be used in the realism-based model, and the Graph Specification references a subsection of these terms. The Graph Specification Validation Service enforces that:

1. All subjects, predicates, and objects referenced by Connection Recipes in the Graph Specification are defined in the application ontology.
2. If domains and/or ranges for predicates are defined in the application ontology, all subjects referenced by Connection Recipes are within the domain, and all objects referenced by Connection Recipes are within the range.
3. All terms in the Graph Specification are defined either in the application ontology or in the Semantic Engine Language ontology.

##### 3.7.2 Transformation Instruction Set Validation Service

The Transformation Instruction Set must be consistent with the Graph Specification’s stated requirements for when a given Connection Recipe must be executed, and all of its terms must be defined. The Transformation Instruction Set Validation Service enforces that:

1. If a Connection Recipe referenced by an Update Specification as an output refers to a subject or object which is referenced by another Connection Recipe in the Graph Specification, and that Connection Recipe must execute if the subject or object exists, then that Connection Recipe must be the output of at least one Update Specification in the Transformation Instruction Set. For example, given the following, an error would be thrown:
  – An Update Specification has as an output a Connection Recipe with object **:patientRole**
  – The Graph Specification contains another Connection Recipe called **:encounterRealizesPatientRole** with subject **:encounter** and object **:patientRole**, which must execute if **:objectExists**
  – No Update Specification has as an output **:encounterRealizesPatientRole**
  – This is an error because an instance of **:patientRole** is being created and its required relationship with an instance of **:encounter** is not being created.
2. All terms in the Transformation Instruction Set must have a type.
3. All terms in the Transformation Instruction Set that are not subjects, predicates, or objects referenced by a Connection Recipe must be defined in the Semantic Engine Language ontology.

##### 3.7.3 Conflict Detection Validation Service

The Conflict Detection Validation Service operates over both the Transformation Instruction Set and Graph Specification. Its goal is to find single Connection Recipes or a set of Connection Recipes that would introduce an inconsistency, and alert the user before the transformation begins. The Conflict Detection Service enforces that:

1. A Connection Recipe’s stated type corresponds with the subject and/or object that it references. For example, if a recipe is an Instance-to-Instance recipe, and the recipe references an object that is a literal value, this would be flagged.
2. Multiple Connection Recipes do not define cardinalities that are mutually inconsistent. A simple example of an inconsistent cardinality loop is shown in **Figure 7**.

**Figure 7:**
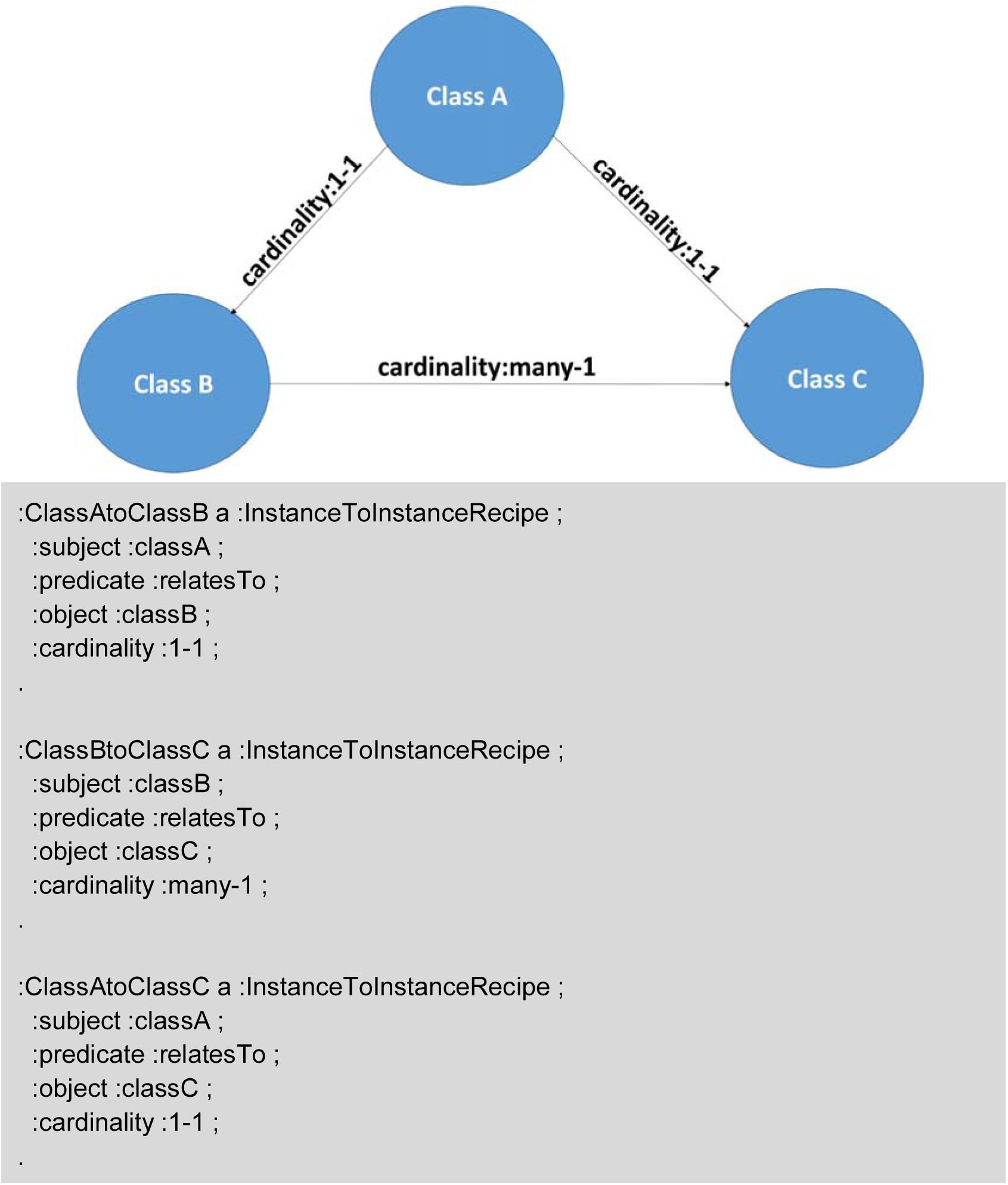
Graphical display and triples of an example of inconsistent cardinality relationships between Connection Recipes in a Transformation Instruction Set

#### 3.7.4 Input Data Validation Service

The Input Data Validation Service is the only Validation Service that operates over the source data that a user has loaded into a repository. It is important that this service run last, because the assumption is that by this point the Semantic Engine Language configuration files have been validated. The input Data Validation Service enforces that:

1. If a Connection Recipe referenced by an Update Specification as a required input must execute on some condition, that condition must be present in the instance data. For example, given the following, an error would be thrown:
  – An Update Specification requires as an input Connection Recipe with subject **inputData:medication and** object **inputData:medOrderName**, which must execute if **:subjectExists**
  – An instance of class **inputData:medication** without an associated instance of **inputData:medOrderName** in the input data exists

#### 3.8 Use Cases

At the point of writing this paper, we have published Transformation Instruction Sets for two data sources. These are openly available on our Github page at https://github.com/PennTURBO/Drivetrain. These Transformation Instruction Sets were created with the intention of instantiating two patient cohorts into our semantic database:

- 51,031 patients with samples in the Penn Medicine Biobank
- 1,153 synthetic patients generated by the Synthea patient data generator^36^

We worked on these Transformation Instruction Sets simultaneously in order to demonstrate the viability of multiple contributors with different data sources working from the same Graph Specification file. This process ensured that similar types of data from each heterogeneous source would be represented identically in the triplestore, thereby achieving interoperability between the data sources.

We built our realism-based model inside our canonical Graph Specification file before beginning work on the Transformation Instruction Sets, keeping our use cases in mind as we chose which data types to include. **Table 4** shows the data fields that our realism-based model includes. Each of these fields is found in either the Penn Medicine Biobank cohort, the Synthea cohort, or both.

**Table 4:** Data fields currently included in the PennTURBO Group’s realism-based model

| Category of Data | Fields Modeled |
| --- | --- |
| Patient demographics and observations | • Centrally Registered Identifier (CRID)<br>• Date of Birth |
|  | • Gender Identity<br>• Racial Identity<br>• Height<br>• Weight<br>• BMI<br>• Systolic Blood Pressure<br>• Diastolic Blood Pressure |
| Healthcare Encounters and Encounters with Biobank Personnel | • Encounter Primary Key<br>• Date of Encounter |
| Diagnoses | • Diagnosis Primary Key<br>• Diagnosis Code<br>• Diagnosis Code Registry (ICD9, ICD10, SNOMED, etc.)<br>• Diagnosis Description String |
| Medications | • Medication Primary Key<br>• Medication Code<br>• Medication Code Registry (RxNorm, CVX, etc.)<br>• Medication Description String |
| Loss of Function Predictions about Genes | • Gene Identifier<br>• Heterozygous or homozygous Loss of Function Mutants (zygosity) |

##### 3.8.1 Penn Medicine Biobank Pipeline Architecture

One of the goals of the PennTURBO project at the University of Pennsylvania is to provide querying services over cohorts of patient data of interest to researchers. One particular cohort of interest is data representing the patients in the Penn Medicine Biobank. Of the roughly 51,000 patients who have donated blood and/or tissue samples that are stored in the Biobank, about 10,000 have given samples that have undergone genomic analysis. Predictions of mutations causing loss-of-function in a particular gene within this cohort are stored in a large CSV file. The rest of the data for this cohort exists in Penn Data Store (PDS), a large clinical data warehouse in the form of an Oracle relational database.

We developed a pipeline to convert the Penn Medicine Biobank data into RDF. First, the Carnival application was able to pull patient demographic information for all Biobank patients from PDS into a property graph adhering to a schema inspired by OBO Foundry ontologies. Then, SQL queries launched from Carnival pulled data about diagnoses, prescriptions, and measurements associated with this patient cohort directly from PDS into application memory. The next step was to pull the loss-of-function data from the CSV file into a temporary database. Finally, we processed the property graph data, the data held in-memory from PDS, and the genomic data within our temporary database, to create a set of concise triples. **Figure 8** shows a visual diagram representing the entire pipeline architecture.

**Figure 8:**
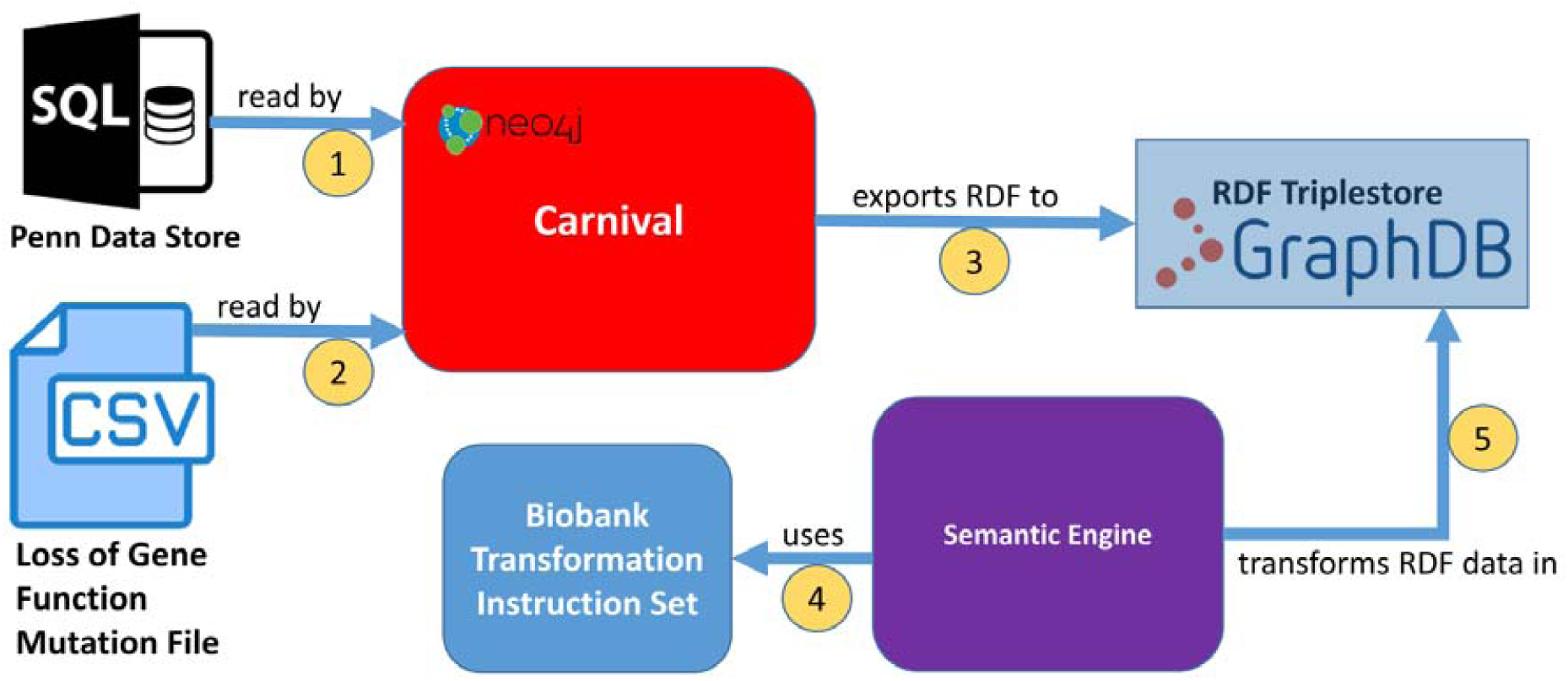
Visual display of the PennMedicine Biobank Pipeline

##### 3.8.2 Synthea Pipeline Architecture

The second Transformation Instruction Set is based on the Observational Medical Outcomes Partnership (OMOP) relational schema. The goals for this effort were 1) prepare for future instantiations of datasets from various sources pre-formatted using the OMOP schema and 2) demonstrate the viability of including multiple data sources simultaneously using the same Graph Specification.

We developed a pipeline to convert the Synthea data into RDF. First, we used Synthea^37^, a synthetic patient data simulator, to generate synthetic data for a roughly 1,000 patient population, including the fields most commonly requested by researchers. Then, the Synthea-generated CSV files were loaded into a PostgreSQL instance. The following step was to use the ETL-Synthea conversion tool^38^, published by Observational Health Data Sciences and Informatics (OHDSI), to migrate the data into new PostgreSQL schemas in the OMOP format. At the time of the migration, the conversion tool used a recent OMOP schema version (5.x). Finally, in order to demonstrate that our pipeline is not dependent on a particular data aggregation service, we chose to perform the OMOP transformation to RDF with Stardog’s Virtual Graph^39^. This service instantiated the Synthea data in the OMOP PostgreSQL database as concise triples. **Figure 9** shows a visual diagram representing the Synthea pipeline architecture.

**Figure 9:**
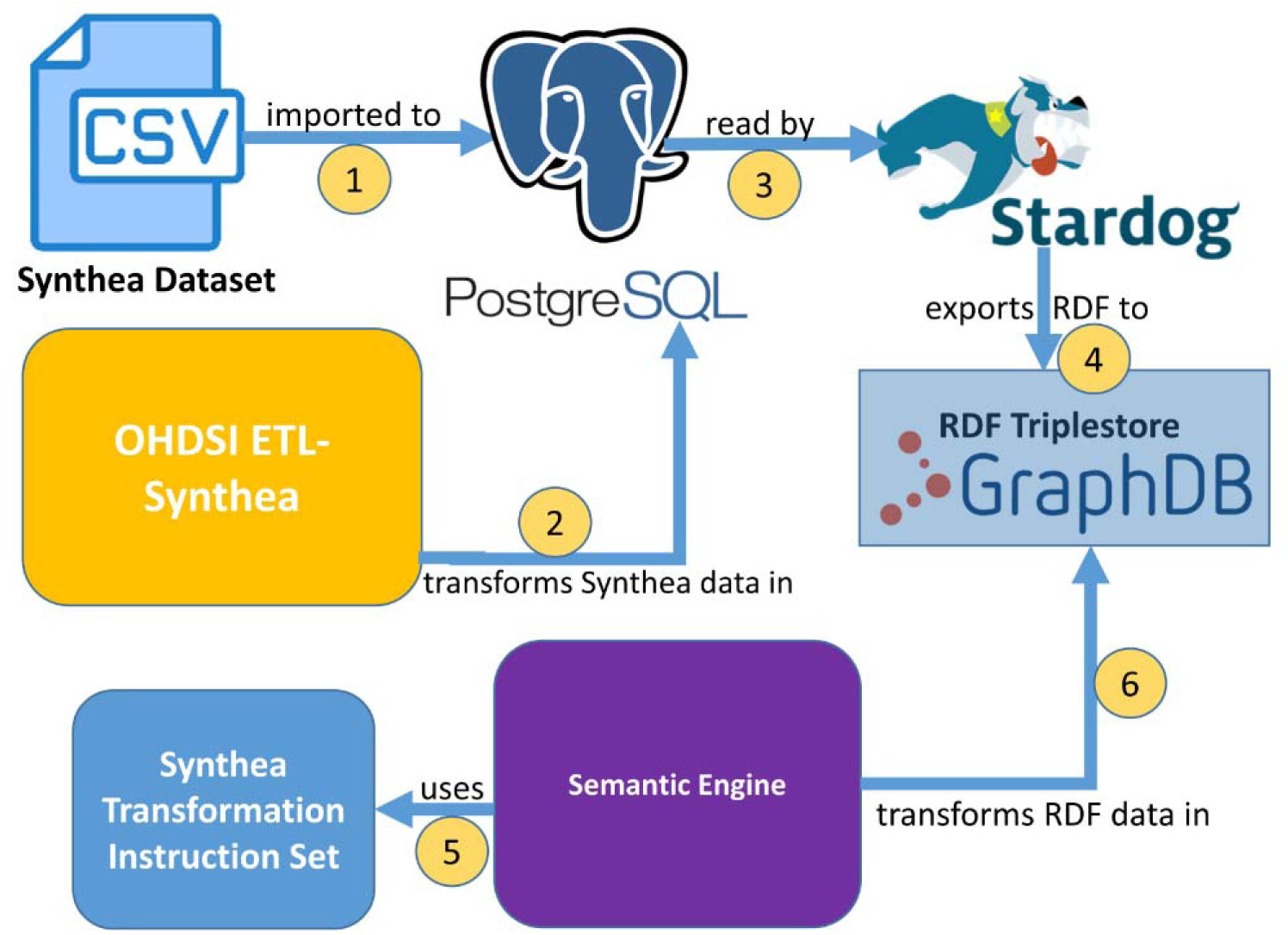
Visual display of the Synthea Pipeline

##### 3.8.3 Transformation Instruction Set Comparison

**Table 5** provides a breakdown of Connection Recipes by type and origin for both the Synthea and Biobank use cases. Recipes originating in the Transformation Instruction Set describe the relationships present in the incoming data source, and Recipes referenced from the Graph Specification represent relationships in the realism-based model. Many of the differences in counts between the Instruction Sets are due to differences in style between the developers of each pipeline.

**Table 5:** Connection Recipe type counts referenced by Update Specifications in Biobank and Synthea Transformation Instruction Sets

| Origin | Instance-to-Instance | Instance-to-Term | Term-to-Instance | Instance-to-Literal | Term-to-Literal | Term-to-Term |
| --- | --- | --- | --- | --- | --- | --- |
| Biobank Instruction Set | 16 | 14 | 0 | 21 | 0 | 0 |
| Biobank Instruction Set Reference to Graph Specification | 50 | 16 | 2 | 21 | 0 | 0 |
| Synthea Instruction Set | 12 | 6 | 0 | 45 | 2 | 2 |
| Synthea Instruction Set Reference to Graph Specification | 117 | 69 | 0 | 36 | 0 | 0 |

One such difference in style is reflected in the presence of Term-to-Literal and Term-to-Term Recipes in the Synthea Instruction Set, but not in the Biobank Instruction Set. In some cases, mappings of concept codes to classes may be required to properly transform a set of input data. For instance, if an incoming data source represents biological sex by a string “M” for male or “F” for female, it would be appropriate for the application ontology classes that represent biological sex concepts to refer to those strings. We tried out two approaches for searching within the application ontology for these mappings:

1. Instruct the Semantic Engine to use a combination of Term-to-Term and Term-to-Instance Recipes to perform the lookup, such as occurs in the Synthea Instruction Set
2. Use an external tool to perform the lookup and include the mappings as part of the input triples, such as occurs in the Biobank Instruction Set, which makes including Term-to-Term Recipes and Term-to-Instance Recipes in that Transformation Instruction Set unnecessary because the relevant mappings have already been provided

Another observation from **Table 5** is that the Synthea Instruction Set uses significantly more Connection Recipes from the Graph Specification than does the Biobank Instruction Set. Our canonical Graph Specification contains many Connection Recipes that record provenance of data from a relational source, generating statements about the specific columns, tables, schemas, and databases from which each data field originated. The developer of the Synthea Instruction Set chose to include statements about provenance of the data, whereas the developer of the Biobank Instruction Set did not. Both solutions generate valid implementations of the realism-based model, since the data provenance Recipes are optional.

#### 3.9 Performance

In measuring the performance of the Semantic Engine running against a dataset, the key factors are the efficiency of the generated SPARQL updates and the resources available to the database server. The time consumed by the Semantic Engine to generate the SPARQL updates is dependent on the size of the Transformation Instruction Set, not the instance data. As Transformation Instruction Set files are generally small, this process usually takes only a few seconds at most. Running of the generated queries against the database takes the bulk of the time.

Our triplestore is Ontotext GraphDB^40^ Standard Edition Version 9.1.1 running on a Linux Kernel using the CentOS Linux distribution, version 6.9. The Standard Edition requires a license but allows queries to run in parallel, which significantly improves the transformation time. We have also tested our system running queries in series using the Free Edition, which is more time consuming but produces the same results. In theory, the Semantic Engine should run with any triplestore that supports RDF4J; however, some modifications to the code that creates the connection to the triplestore might be necessary.

To effectively benchmark our system at various throughput levels, we used Synthea to generate a dataset of one million synthetic patients, including their demographics, measurements, medications, and diagnoses, and stored this data in a PostgreSQL database. We then used the ETL-Synthea tool to format the data to conform to the OMOP schema.^41^ This transformation was helpful because Carnival already included facilities for reading and transforming OMOP-formatted data. Finally, we used Carnival to format the synthetic OMOP data into RDF.

Once the pipeline was completed, our final steps were to generate concise RDF datasets of various sizes, which we controlled by limiting the number of patients included, and then to run the Semantic Engine against these datasets. The charts below demonstrate the Carnival application’s performance for building the source-specific RDF dataset from the data in the PostgreSQL database, as well as the Semantic Engine’s performance transforming the source-specific RDF dataset. We ran cohorts of 1,000, 10,000, 100,000, and 1,000,000 patients. **Table 6** shows the types and quantities of data included in each of these cohorts.

**Table 6:** Size and counts of entities in Synthea cohorts generated for performance testing

| Number of Initial Triples from Carnival | Number of Final Triples after Semantic Engine run | Counts of Included Entities |
| --- | --- | --- |
| 917,369 | 3,593,773 | • 1,000 patients<br>• 36,184 healthcare encounters<br>• 6,727 diagnoses<br>• 46,252 prescriptions<br>• 51,163 measurements |
| 9,380,085 | 36,609,719 | • 10,000 patients<br>• 372,297 healthcare encounters<br>• 69,484 diagnoses<br>• 482,093 prescriptions<br>• 507,875 measurements |
| 89,808,702 | 350,150,937 | • 100,000 patients<br>• 3,587,004 healthcare encounters<br>• 675,998 diagnoses<br>• 4,516,160 prescriptions<br>• 4,917,038 measurements |
| 894,416,550 | 3,487,368,869 | • 1,000,000 patients |
|  |  | • 35,786,086 healthcare encounters<br>• 6,703,298 diagnoses<br>• 45,073,629 prescriptions<br>• 48,793,260 measurements |

In **Figure 10**, we see a comparison of the time needed for Carnival to instantiate the initial RDF triples, and for the Semantic Expander to transform those triples into the realism-based model. Carnival uses a batched process to export the triples that involves clearing its embedded property graph after each chunk of data is pushed to the triplestore and before a new chunk is pulled from the PostgreSQL database. This hurts performance at low throughput levels but ensures that it can scale linearly as throughput levels increase.

**Figure 10:**
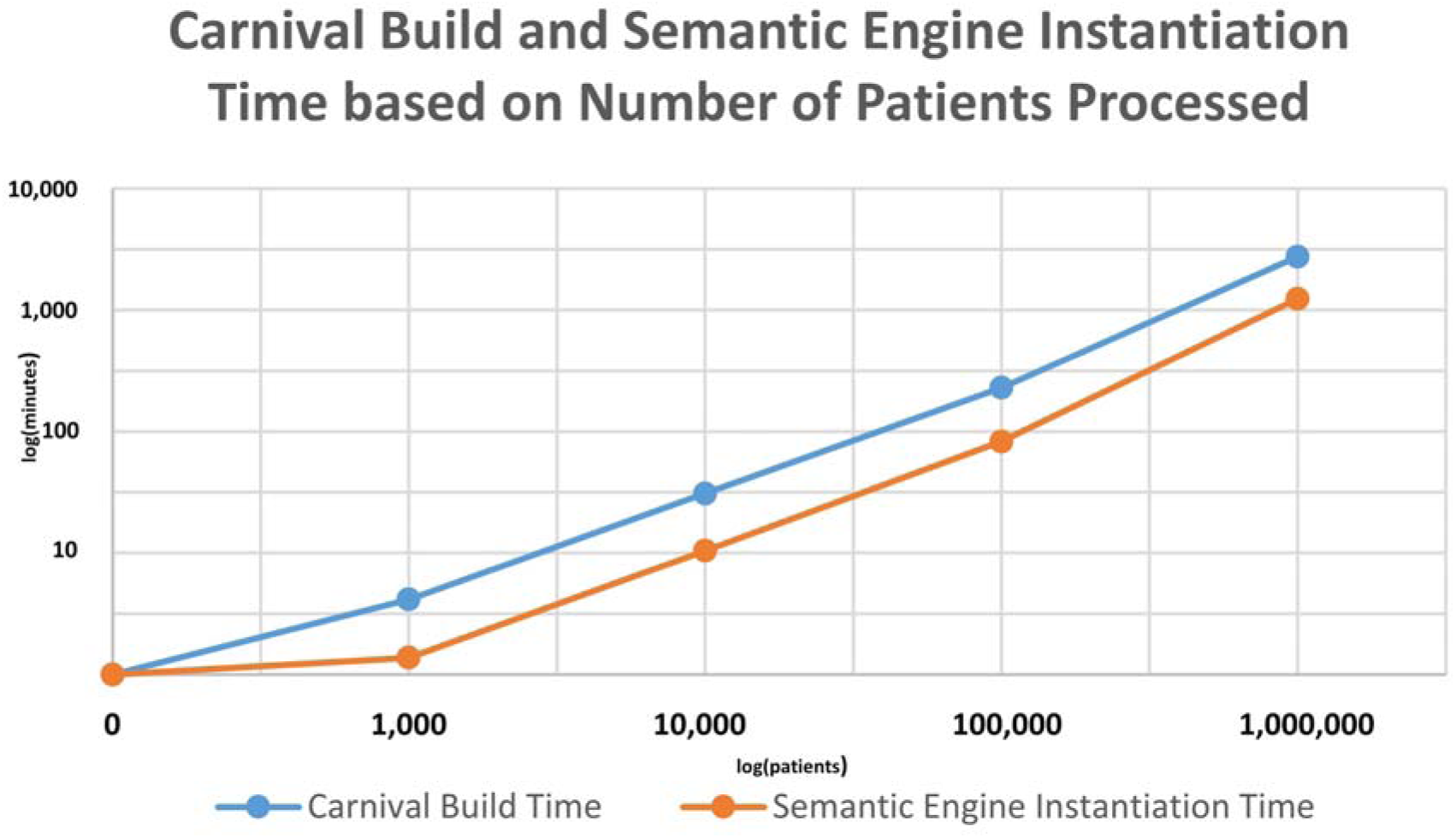
Comparison of the time taken for Carnival to build source-specific RDF dataset, and to instantiate it using the Semantic Expander, by cohort size. The graph shows an approximately linear relationship between size and time.

In **Figure 11**, we see that the performance of the Semantic Expander was mostly linear over the specified throughput levels for each type of data. This is encouraging, as we plan to explore the potential of our system for enhancing data from large data lakes and warehouses.

**Figure 11:**
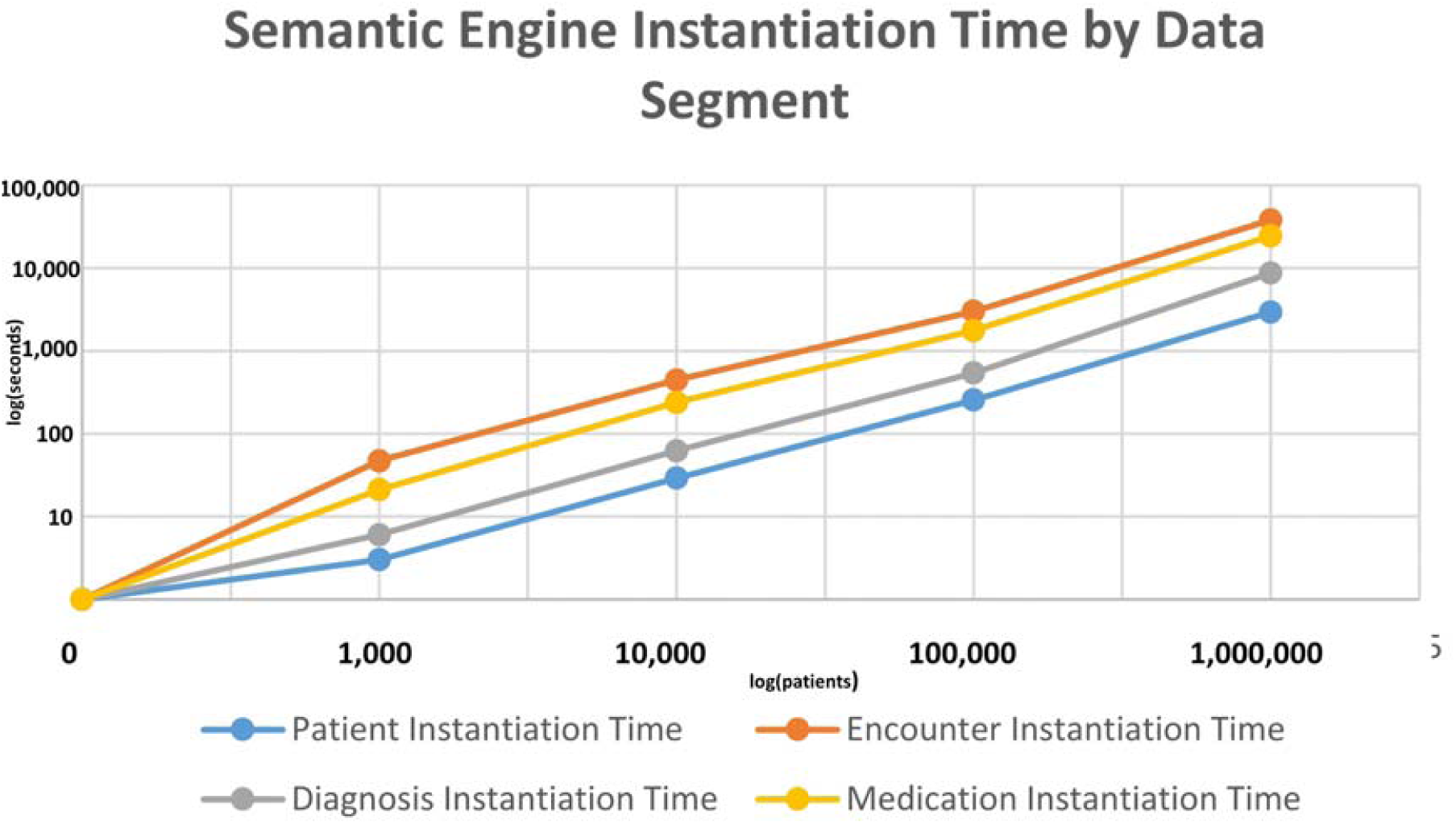
Comparison of the time taken for the Semantic Expander to transform various types of data, by cohort size. The graph shows an approximately linear relationship between size and time.

## 4. Discussion

### 4.1 Generalizability

Our development of the Semantic Engine was guided by principles of generalizability. Although we conceived of the application with a specific use case in mind, we ensured that any use case-specific elements were specified in components external to the software code itself, such as our application ontology, Graph Specification, and Transformation Instruction Sets. These components can be swapped in and out as needed. We anticipate different ways that outside groups could take advantage of the work we have done, based on their domains of interest:

- For clinical data projects involving the same or similar fields we have already modeled, applying our vetted model to native data sources by crafting their own Transformation Instruction Sets, and using our canonical Graph Specification and the PennTURBO application ontology
- For clinical data projects necessitating the inclusion of fields our group has not already modeled, adding the new fields to our vetted model by modifying the Graph Specification and then applying it to native data sources by crafting their own Transformation Instruction Sets, and using the amended version of the Graph Specification and the PennTURBO application ontology
- For projects interested in domains outside of the clinical data sphere, creating their own application ontology and data model based on their project’s needs, then applying it to native data sources by crafting their own Graph Specification and Transformation Instruction Sets
- For clinical or non-clinical projects which already have an application ontology which suits their needs, building a Graph Specification and Transformation Instruction Sets to go along with the pre-existing ontology. The Semantic Engine was initially specified for realism-based ontology models; however, a non-OBO Foundry-compliant application ontology could be used as well.

We feel that the principles that guided our construction of the Semantic Engine allow it to work with nearly any domain or data source. However, the process of creating a Semantic Engine configuration can be time consuming and may require the input of domain experts. Additionally, certain more complex use cases might require the implementation of several advanced language features not discussed in this paper. A future area of work might be to create software tools to assist in the building of Semantic Engine Language configurations.

### 4.2 Limitations

One of the shortcomings of the Semantic Engine is that it is limited by the generic RDF data model, the SPARQL language, and triplestore implementations. Given an especially large input dataset or complex specification, a query could be generated which takes a large amount of time to complete. We recommend implementing multiple SPARQL queries for the transformation of large input datasets. Some triplestores have a Query Explain service that can also be helpful in determining the complexity of a query before it is executed. Initially trying a pipeline on a small dataset can be helpful for gauging performance of the full dataset, since we have observed performance to scale linearly with the size of the data. We have also noticed queries are more likely to complete without crashing when we set an infinite (or very high) timeout setting for transactions on our triplestore system.

An area for future improvement of the Semantic Engine Language is the ability to express more complicated transformation rules for literal values. Examples of operations users may wish to define include concatenating two strings, selecting an ontology class based on the value of a literal, or constructing a URI that includes the literal value to create a pointer to an external terminology. We currently implement these special cases by allowing the user to submit snippets of custom SPARQL in the Transformation Instruction Set; however, even in our preliminary use of the tool we have seen that this introduces a potential source of errors. The solution proposed by Mate et. al handles this situation nicely by implementing a suite of functions such as “ADD” which perform basic functions on literal values coming from their data source.^16^

The Semantic Engine Language provides a custom solution for the problem of specifying RDF graph schemas; however especially in the context of graph validation, there are some pre-existing options. As of its latest release, GraphDB now supports SHACL, which provides a syntax for creating RDF shapes and using them to validate RDF instance data. We may continue to explore the use of SHACL as a potential replacement for or addendum to the validation already provided by the Semantic Engine. Additionally, it might be interesting to create a framework for translating Semantic Engine Language configuration files into SHACL, and vice versa.

## 5. Conclusion

The ability to respond quickly and accurately to investigators with granular clinical data requests is important to facilitate high quality, impactful research. We have presented a system that leverages OBO Foundry ontologies to construct a realism-based model for storing and querying clinical data, and that has the ability to transform domain-specific data using declaratively specified, dynamically generated SPARQL update statements. Our realism-based model has been highly vetted by members of the OBI and OBIB communities and is inherently documented and validated by the Semantic Engine Language, allowing it to be shared with and applied by data brokers at other institutions. By modeling our data with a high level of semantic richness, we abide by one of the original principles of semantic Web technologies: that well-defined data can unlock major new functionality.^42^

We have reviewed similar efforts to create clinical data models with high potential for interoperability. Sun et al. and Berges et al. similarly first shape their data into source-specific RDF and then transform the data to conform to target models. Both use well-known reasoner engines (EYE and SWRL, respectively) to complete the transformation. Our solution to use a pure SPARQL solution executed by a custom DSL was advantageous in comparison to the solutions we surveyed in that it 1) separates transformation instructions from the target model, 2) is easily shareable, and 3) allows for automatic validation of a user’s transformation instructions and instance data.

We describe the syntax of the Semantic Engine Language and show examples of how it compiles to SPARQL. Since SPARQL provides the underlying implementation for the Semantic Engine, the system can leverage its fast performance and RDF-specific features without forcing a user to handcraft long, error-prone queries. Because a user is insulated from the query that performs their process, they can focus their energy on crafting the target model itself. Rather than hand checking that they have consistently implemented their transformation within a single query or between multiple queries running in sequence, they can leverage the Semantic Engine’s ability to re-use patterns and self-validate to ensure consistency.

We believe the Semantic Engine can facilitate interdisciplinary and inter-institutional collaboration towards the creation of a widely accepted realism-based model for representing patient data. The Semantic Engine is currently used at the University of Pennsylvania as part of the PennTURBO project, an effort to consolidate relational data about patients from disparate sources into cohorts of interest, represented in RDF. We encourage published additions to our canonical Graph Specification to increase the scope of the types of data that are modeled.

Our open-source code base as well as our Semantic Engine configuration files are available at https://github.com/PennTURBO/Drivetrain.

## Acknowledgements

This work was done as part of the PennTURBO project, which is supported by the Institute for Biomedical Informatics and by the Institute for Translational Medicine and Therapeutics at the University of Pennsylvania.

We would like to thank Alan H. Ruttenberg, Bill Duncan, and James A. Overton for their constructive comments on an early version of the manuscript.

## Notes

### Competing Interest Statement

The authors have declared no competing interest.

